## Supplementary material for "Switching state-space modeling of neural signal dynamics": S1 Appendix. Free energy calculation

### Computation of negative free energy

To compute the negative variational free energy $\mathcal{F}$, we start from its definition:

$$\begin{aligned}\mathcal{F}(q,\boldsymbol{\theta}) &= \mathbb{E}_q\left[\log\frac{p\left(\left\{\mathbf{y}_t,\mathbf{x}_t^{(1)},\cdots,\mathbf{x}_t^{(M)},s_t\right\}|\boldsymbol{\theta}\right)}{q\left(\left\{\mathbf{x}_t^{(1)},\cdots,\mathbf{x}_t^{(M)},s_t\right\}\right)}\right]\\ &= \left\langle\left\langle\log p\left(\left\{\mathbf{y}_t,\mathbf{x}_t^{(1)},\cdots,\mathbf{x}_t^{(M)},s_t\right\}|\boldsymbol{\theta}\right)-\log q\left(\{s_t\}\right)-\log q\left(\left\{\mathbf{x}_t^{(1)},\cdots,\mathbf{x}_t^{(M)}\right\}\right)\right\rangle_s\right\rangle_x \quad (1)\end{aligned}$$

In Materials and methods, we have developed the log-posterior expression for the switching state:

$$\begin{aligned}\log q\left(\{s_t\}\right) &= \left\langle\log p\left(\left\{\mathbf{y}_t,\mathbf{x}_t^{(1)},\cdots,\mathbf{x}_t^{(M)},s_t\right\}|\boldsymbol{\theta}\right)\right\rangle_x-\log z\\ &= \sum_{m=1}^{M}\mathbb{1}[s_0=m]\log\boldsymbol{\rho}_m+\sum_{t=1}^{T}\sum_{m,n=1}^{M}\mathbb{1}\left[s_t=m,s_{t-1}=n\right]\log\boldsymbol{\phi}_{m,n}\\ &\quad+\sum_{t=1}^{T}\sum_{m=1}^{M}\mathbb{1}[s_t=m]g_t^{(m)}-\log\zeta \quad (2)\\ &\quad\text{with } g_t^{(m)}\triangleq-\frac{1}{2}\left\langle\left(\mathbf{y}_t-\mathbf{G}^{(m)}\mathbf{x}_t^{(m)}\right)^\top\mathbf{R}^{-1}\left(\mathbf{y}_t-\mathbf{G}^{(m)}\mathbf{x}_t^{(m)}\right)\right\rangle_x\end{aligned}$$

and similarly for the real-valued Gaussian SSMs:

$$\begin{aligned}\log q\left(\left\{\mathbf{x}_t^{(1)},\cdots,\mathbf{x}_t^{(M)}\right\}\right) &= \left\langle\log p\left(\left\{\mathbf{y}_t,\mathbf{x}_t^{(1)},\cdots,\mathbf{x}_t^{(M)},s_t\right\}|\boldsymbol{\theta}\right)\right\rangle_s-\log z'\\ &= -\frac{1}{2}\sum_{m=1}^{M}\left(\mathbf{x}_0^{(m)}-\boldsymbol{\mu}^{(m)}\right)^\top{\mathbf{Q}_0^{(m)}}^{-1}\left(\mathbf{x}_0^{(m)}-\boldsymbol{\mu}^{(m)}\right)\\ &\quad-\frac{1}{2}\sum_{m=1}^{M}\sum_{t=1}^{T}\left(\mathbf{x}_t^{(m)}-\mathbf{F}^{(m)}\mathbf{x}_{t-1}^{(m)}\right)^\top{\mathbf{Q}^{(m)}}^{-1}\left(\mathbf{x}_t^{(m)}-\mathbf{F}^{(m)}\mathbf{x}_{t-1}^{(m)}\right)\\ &\quad-\frac{1}{2}\sum_{t=1}^{T}\sum_{m=1}^{M}h_t^{(m)}\left(\mathbf{y}_t-\mathbf{G}^{(m)}\mathbf{x}_t^{(m)}\right)^\top\mathbf{R}^{-1}\left(\mathbf{y}_t-\mathbf{G}^{(m)}\mathbf{x}_t^{(m)}\right)-\log\zeta' \quad (3)\\ &\quad\text{with } h_t^{(m)}\triangleq\left\langle\mathbb{1}[s_t=m]\right\rangle_s\end{aligned}$$

Recall that the complete log-likelihood expression is:

$$\begin{aligned}
&\log p\left(\left\{\mathbf{y}_t, \mathbf{x}_t^{(1)}, \cdots, \mathbf{x}_t^{(M)}, s_t\right\} |\boldsymbol{\theta}\right) \\
&= -\frac{T}{2}\log|2\pi\mathbf{R}| - \frac{1}{2}\sum_{t=1}^{T}\sum_{m=1}^{M}\mathbb{1}[s_t=m]\left(\mathbf{y}_t - \mathbf{G}^{(m)}\mathbf{x}_t^{(m)}\right)^\top \mathbf{R}^{-1}\left(\mathbf{y}_t - \mathbf{G}^{(m)}\mathbf{x}_t^{(m)}\right) \\
&\quad - \frac{1}{2}\sum_{m=1}^{M}\log\left|2\pi\mathbf{Q}_0^{(m)}\right| - \frac{1}{2}\sum_{m=1}^{M}\left(\mathbf{x}_0^{(m)} - \boldsymbol{\mu}^{(m)}\right)^\top \mathbf{Q}_0^{(m)-1}\left(\mathbf{x}_0^{(m)} - \boldsymbol{\mu}^{(m)}\right) \\
&\quad - \frac{T}{2}\sum_{m=1}^{M}\log\left|2\pi\mathbf{Q}^{(m)}\right| - \frac{1}{2}\sum_{m=1}^{M}\sum_{t=1}^{T}\left(\mathbf{x}_t^{(m)} - \mathbf{F}^{(m)}\mathbf{x}_{t-1}^{(m)}\right)^\top \mathbf{Q}^{(m)-1}\left(\mathbf{x}_t^{(m)} - \mathbf{F}^{(m)}\mathbf{x}_{t-1}^{(m)}\right) \\
&\quad + \sum_{m=1}^{M}\mathbb{1}[s_0=m]\log\boldsymbol{\rho}_m + \sum_{t=1}^{T}\sum_{m,n=1}^{M}\mathbb{1}\left[s_t=m, s_{t-1}=n\right]\log\boldsymbol{\phi}_{m,n}
\end{aligned} \tag{4}$$

Therefore, most of the terms in Eq 1 cancel out except the log normalization constants $\log\zeta$ and $\log\zeta^{'}$ along with some determinant constants in Eq 4 that are currently absorbed in $\log\zeta^{'}$ in Eq 3. It is clear that we now need to efficiently compute the normalization constants $\zeta$ and $\zeta^{'}$.

To compute $\zeta$, observe that Eq 2 is already in the same form of a hidden Markov model process:

$$p\left(\{\mathbf{y}_t, s_t\}\right) = p\left(s_0\right)\prod_{t=1}^{T} p\left(s_t|s_{t-1}\right) p\left(\mathbf{y}_t|s_t\right)$$

We can then directly use the results from [1, Section III-A, *The Forward-Backward Procudure*]:

$$1 = \sum_{m=1}^{M} q\left(\{s_t = m\}\right) = \frac{1}{\zeta}\sum_{m=1}^{M}\alpha_T(m)$$

Therefore,

$$\zeta = \sum_{m=1}^{M}\alpha_T(m) \tag{5}$$

where $\alpha_T(m)$ is the forward pass variable at the last time point in the forward-backward algorithm that has already been completed as part of the fixed-point iterations in the E-step. Note that because we typically have long recordings with a high sampling rate, using this definition directly is computationally unstable. We prove a normalized version of Eq 5 in S2 Appendix.

To compute $\zeta^{'}$, we would hope to apply the same technique and rewrite Eq 3 into the same form of a set of Gaussian SSMs on which we have completed Kalman filtering:

$$p\left(\left\{\mathbf{y}_t, \mathbf{x}_t^{(1)}, \cdots, \mathbf{x}_t^{(M)}\right\}\right) = \prod_{m=1}^{M}\left(p\left(\mathbf{x}_0^{(m)}\right)\prod_{t=1}^{T} p\left(\mathbf{x}_t^{(m)}|\mathbf{x}_{t-1}^{(m)}\right) p\left(\mathbf{y}_t|\mathbf{x}_t^{(m)}\right)\right)$$

One complication is that due to the last term before the normalization constant in Eq 3, Kalman filtering and smoothing were carried out using $\mathbf{R}/h_t^{(m)}$ as the observation noise covariance. The corresponding complete log-likelihood function for such set of Gaussian SSMs and observations is:

$$\begin{aligned}
&\log\, p\left(\left\{\mathbf{y}_t, \mathbf{x}_t^{(1)}, \cdots, \mathbf{x}_t^{(M)}\right\}|\boldsymbol{\theta}\right) \\
&= -\frac{1}{2}\sum_{m=1}^{M}\sum_{t=1}^{T}\log\left|2\pi\left(\frac{\mathbf{R}}{h_t^{(m)}}\right)\right| - \frac{1}{2}\sum_{m=1}^{M}\sum_{t=1}^{T}\left(\mathbf{y}_t - \mathbf{G}^{(m)}\mathbf{x}_t^{(m)}\right)^{\top}\left(\frac{\mathbf{R}}{h_t^{(m)}}\right)^{-1}\left(\mathbf{y}_t - \mathbf{G}^{(m)}\mathbf{x}_t^{(m)}\right) \\
&\quad - \frac{1}{2}\sum_{m=1}^{M}\log\left|2\pi\mathbf{Q}_0^{(m)}\right| - \frac{1}{2}\sum_{m=1}^{M}\left(\mathbf{x}_0^{(m)} - \boldsymbol{\mu}^{(m)}\right)^{\top}\mathbf{Q}_0^{(m)^{-1}}\left(\mathbf{x}_0^{(m)} - \boldsymbol{\mu}^{(m)}\right) \\
&\quad - \frac{T}{2}\sum_{m=1}^{M}\log\left|2\pi\mathbf{Q}^{(m)}\right| - \frac{1}{2}\sum_{m=1}^{M}\sum_{t=1}^{T}\left(\mathbf{x}_t^{(m)} - \mathbf{F}^{(m)}\mathbf{x}_{t-1}^{(m)}\right)^{\top}\mathbf{Q}^{(m)^{-1}}\left(\mathbf{x}_t^{(m)} - \mathbf{F}^{(m)}\mathbf{x}_{t-1}^{(m)}\right) \qquad (6)
\end{aligned}$$

Since Eq 3 and Eq 6 only differ in constants, we slightly abuse the $\zeta^{'}$ notation and re-write Eq 3 as:

$$\log q\left(\left\{\mathbf{x}_t^{(1)}, \cdots, \mathbf{x}_t^{(M)}\right\}\right) = \log p\left(\left\{\mathbf{y}_t, \mathbf{x}_t^{(1)}, \cdots, \mathbf{x}_t^{(M)}\right\}|\boldsymbol{\theta}\right) - \log\zeta^{'} \qquad (7)$$

Similar to Eq 2, the first term on the right-hand side of Eq 7 now has the same form of Gaussian SSM densities that can be integrated efficiently, allowing us to write:

$$\begin{aligned}
1 &= \int q\left(\left\{\mathbf{x}_t^{(1)}, \cdots, \mathbf{x}_t^{(M)}\right\}\right)\, d\left\{\mathbf{x}_t^{(1)}, \cdots, \mathbf{x}_t^{(M)}\right\} \\
&= \frac{1}{\zeta^{'}}\int p\left(\left\{\mathbf{y}_t, \mathbf{x}_t^{(1)}, \cdots, \mathbf{x}_t^{(M)}\right\}|\boldsymbol{\theta}\right)\, d\left\{\mathbf{x}_t^{(1)}, \cdots, \mathbf{x}_t^{(M)}\right\} = \frac{1}{\zeta^{'}}p\left(\left\{\mathbf{y}_t\right\}|\boldsymbol{\theta}\right)
\end{aligned}$$

Therefore,

$$\zeta^{'} = p\left(\left\{\mathbf{y}_t\right\}|\boldsymbol{\theta}\right) \qquad (8)$$

where $p(\{\mathbf{y}_t\}|\boldsymbol{\theta})$ is simply the product of marginal log-likelihoods of observations in the set of $M$ Gaussian SSMs during Kalman filtering. We can use the results from [2] to compute $\log p(\{\mathbf{y}_t\}|\boldsymbol{\theta})$ in the innovations form:

$$\begin{aligned}
\log p\left(\left\{\mathbf{y}_t\right\}|\boldsymbol{\theta}\right) = &-\frac{1}{2}\sum_{m=1}^{M}\sum_{t=1}^{T}\log\left|2\pi\left(\mathbf{G}^{(m)}\boldsymbol{\Sigma}_{t|t-1}^{(m)}\mathbf{G}^{(m)^{\top}} + \frac{\mathbf{R}}{h_t^{(m)}}\right)\right| \\
&-\frac{1}{2}\sum_{m=1}^{M}\sum_{t=1}^{T}\left(\mathbf{y}_t - \mathbf{G}^{(m)}\mathbf{x}_{t|t-1}^{(m)}\right)^{\top}\left(\mathbf{G}^{(m)}\boldsymbol{\Sigma}_{t|t-1}^{(m)}\mathbf{G}^{(m)^{\top}} + \frac{\mathbf{R}}{h_t^{(m)}}\right)^{-1}\left(\mathbf{y}_t - \mathbf{G}^{(m)}\mathbf{x}_{t|t-1}^{(m)}\right)
\end{aligned}$$

where $\mathbf{x}_{t|t-1}^{(m)}$ and $\boldsymbol{\Sigma}_{t|t-1}^{(m)}$ are the predicted estimates of hidden states during Kalman filtering that have already been completed as part of the fixed-point iterations in the E-step.

Taken together, substituting Eq 2-8 into Eq 1, we obtain the expression for $\mathcal{F}$:

$$\begin{aligned}
\mathcal{F}(q, \boldsymbol{\theta}) = &-\frac{T}{2}\log\left|2\pi\mathbf{R}\right| + \log\sum_{m=1}^{M}\alpha_T(m) - \sum_{m=1}^{M}\sum_{t=1}^{T}h_t^{(m)}g_t^{(m)} \\
&+\left(\frac{1}{2}\sum_{m=1}^{M}\sum_{t=1}^{T}\log\left|2\pi\left(\frac{\mathbf{R}}{h_t^{(m)}}\right)\right| + \log p\left(\left\{\mathbf{y}_t\right\}|\boldsymbol{\theta}\right)\right)
\end{aligned}$$
