## Supplementary material for "Switching state-space modeling of neural signal dynamics": S2 Appendix. Joint estimation of parameters

### Bivariate AR1 with shared parameters

As described in the section in Results, the true generative model in this simulation example is:

$$\begin{aligned} \mathbf{x}_t &= \begin{bmatrix} 0.5 & \mathbf{F}_{12}^{(s_t)} \\ 0 & 0.5 \end{bmatrix} \mathbf{x}_{t-1} + \mathbf{w}_t, & \mathbf{w}_t &\sim \mathcal{N}\left(\mathbf{0}, \begin{bmatrix} 2 & 0 \\ 0 & 2 \end{bmatrix}\right) \\ \mathbf{y}_t &= \begin{bmatrix} 1 & 0 \\ 0 & 1 \end{bmatrix} \mathbf{x}_t + \mathbf{v}_t, & \mathbf{v}_t &\sim \mathcal{N}\left(\mathbf{0}, \begin{bmatrix} 0.1 & 0 \\ 0 & 0.1 \end{bmatrix}\right) \\ &\text{with} \quad \mathbf{F}_{12}^{(1)} = 0.5, \quad \mathbf{F}_{12}^{(2)} = 0 \end{aligned} \tag{1}$$

By construction, the variational learning algorithm assumes a set of parallel Gaussian SSM to represent any switching dynamic. This means the assumed generative structure is instead:

$$\begin{aligned} \mathbf{x}_t^{(1)} &= \mathbf{F}^{(1)} \mathbf{x}_{t-1}^{(1)} + \mathbf{w}_t^{(1)}, & \mathbf{w}_t^{(1)} &\sim \mathcal{N}\left(\mathbf{0}, \mathbf{Q}^{(1)}\right) \\ \mathbf{x}_t^{(2)} &= \mathbf{F}^{(2)} \mathbf{x}_{t-1}^{(2)} + \mathbf{w}_t^{(2)}, & \mathbf{w}_t^{(2)} &\sim \mathcal{N}\left(\mathbf{0}, \mathbf{Q}^{(2)}\right) \\ \mathbf{y}_t &= \mathbf{G}^{(s_t)} \mathbf{x}_t^{(s_t)} + \mathbf{v}_t, & \mathbf{v}_t &\sim \mathcal{N}\left(\mathbf{0}, \mathbf{R}\right) \end{aligned}$$

where the discrete switching state follows a hidden Markov model (HMM) process with binary states. It is obvious that if we apply the variational method naively, it diverges far from the target generative model in Eq 1. To approximate the desired multi-modality, we can write a hybrid generative model as follows:

$$\mathbf{x}_t^{(1)} = \begin{bmatrix} \mathbf{F}_{11} & \mathbf{F}_{12}^{(1)} \\ \mathbf{F}_{21} & \mathbf{F}_{22} \end{bmatrix} \mathbf{x}_{t-1}^{(1)} + \mathbf{w}_t^{(1)}, \qquad \mathbf{w}_t \sim \mathcal{N}\left(\mathbf{0}, \mathbf{Q}\right) \tag{2}$$

$$\mathbf{x}_t^{(2)} = \begin{bmatrix} \mathbf{F}_{11} & \mathbf{F}_{12}^{(2)} \\ \mathbf{F}_{21} & \mathbf{F}_{22} \end{bmatrix} \mathbf{x}_{t-1}^{(2)} + \mathbf{w}_t^{(2)}, \qquad \mathbf{w}_t \sim \mathcal{N}\left(\mathbf{0}, \mathbf{Q}\right) \tag{3}$$

$$\mathbf{y}_t = \mathbf{G}\mathbf{x}_t + \mathbf{v}_t, \qquad \mathbf{v}_t \sim \mathcal{N}\left(\mathbf{0}, \mathbf{R}\right) \tag{4}$$

where $\mathbf{G}$ is the same across the two candidate models just like $\mathbf{R}$.

In words, we are approximating a process with switching state-transition matrices in Eq 1 using two parallel models whose hidden states could evolve separately. But the parameters are shared across the models based on prior knowledge of the generative process. Any multi-model structure with a finite number of switching states, which is the case in an HMM, can always be recast into a set of parallel models capturing the same switching pattern. In fact, the observation equation in Eq 4 is identical to that in Eq 1. The key difference is that the parallel models in Eq 2 and Eq 3 have separate hidden states $\mathbf{x}_t^{(1)}$ and $\mathbf{x}_t^{(2)}$. Thus in this case, the true posterior being approached during the E-step is itself an approximation to a different graph with distinct support (i.e., different hidden state nodes). Nevertheless, the real-valued hidden states appear in our generalized EM iterations through posteriors that are smoothed estimates based on observations, which in essence "anchor" the hidden states. In addition, the time points when a model is irrelevant are "suppressed" during weighted Kalman smoothing that scales observation noise $\mathbf{R}$ by the model responsibilities $h_t^{(m)}$. Thus, this parallel model approach gives a reasonable approximation to the switching multi-modal Gaussian process, but conforms to an efficient variational learning algorithm described in this paper.

We now derive the corresponding M-step update equations for the shared parameters in Eq 2-4, i.e., joint estimation across switching models. To break the symmetry between the two candidate models, we set $\mathbf{F}_{12}^{(2)} = 0$. Then, similar to the general case, we start with the expectation of the complete log-likelihood w.r.t the approximate distribution $q$:

$$\begin{aligned}
&\mathbb{E}\left[\log p\left(\left\{\mathbf{y}_t, \mathbf{x}_t^{(1)}, \mathbf{x}_t^{(2)}, s_t\right\} | \boldsymbol{\theta}\right)\right] \\
&= -\frac{T}{2}\log|2\pi\mathbf{R}| - \frac{1}{2}\sum_{m=1}^{2}\log|2\pi\mathbf{Q}_0| - \frac{T}{2}\sum_{m=1}^{2}\log|2\pi\mathbf{Q}| \\
&\quad - \frac{1}{2}\sum_{m=1}^{2}\sum_{t=1}^{T} h_t^{(m)} \operatorname{Tr}\left\{\mathbf{R}^{-1}\left(\left(\mathbf{y}_t - \mathbf{G}\mathbf{x}_{t|T}^{(m)}\right)\left(\mathbf{y}_t - \mathbf{G}\mathbf{x}_{t|T}^{(m)}\right)^\top + \mathbf{G}\boldsymbol{\Sigma}_{t|T}^{(m)}\mathbf{G}^\top\right)\right\} \\
&\quad - \frac{1}{2}\sum_{m=1}^{2}\operatorname{Tr}\left\{\mathbf{Q}_0^{-1}\left(\left(\mathbf{x}_{0|T}^{(m)} - \boldsymbol{\mu}\right)\left(\mathbf{x}_{0|T}^{(m)} - \boldsymbol{\mu}\right)^\top + \boldsymbol{\Sigma}_{0|T}^{(m)}\right)\right\} \\
&\quad - \frac{1}{2}\sum_{m=1}^{2}\operatorname{Tr}\left\{\mathbf{Q}^{-1}\left(\mathbf{C}^{(m)} - \mathbf{B}^{(m)}\mathbf{F}^{(m)\top} - \mathbf{F}^{(m)}\mathbf{B}^{(m)\top} + \mathbf{F}^{(m)}\mathbf{A}^{(m)}\mathbf{F}^{(m)\top}\right)\right\} \\
&\quad + \sum_{m=1}^{2} p_{0|T}(m)\log\boldsymbol{\rho}_m + \sum_{t=1}^{T}\sum_{m,n=1}^{2} p_{t,t-1|T}(m,n)\log\boldsymbol{\phi}_{m,n}
\end{aligned} \tag{5}$$

with

$$\mathbf{F}^{(1)} = \begin{bmatrix}\mathbf{F}_{11} & \mathbf{F}_{12}^{(1)} \\ \mathbf{F}_{21} & \mathbf{F}_{22}\end{bmatrix}, \qquad \mathbf{F}^{(2)} = \begin{bmatrix}\mathbf{F}_{11} & 0 \\ \mathbf{F}_{21} & \mathbf{F}_{22}\end{bmatrix}$$

Then the closed-form update equations at the $i$th EM iteration can be obtained by taking individual partial derivatives. In particular, the joint estimation of some shared parameters has an easy and intuitive form:

$$\boldsymbol{\mu}^i = \frac{1}{2}\sum_{m=1}^{2}\mathbf{x}_{0|T}^{(m)} \tag{6}$$

$$\mathbf{Q}_0^i = \frac{1}{2}\sum_{m=1}^{2}\left(\boldsymbol{\Sigma}_{0|T}^{(m)} + \mathbf{x}_{0|T}^{(m)}\mathbf{x}_{0|T}^{(m)\top} - \mathbf{x}_{0|T}^{(m)}\boldsymbol{\mu}^{i\top} - \boldsymbol{\mu}^i\mathbf{x}_{0|T}^{(m)\top} + \boldsymbol{\mu}^i\boldsymbol{\mu}^{i\top}\right) \tag{7}$$

$$\mathbf{Q}^i = \frac{1}{2T}\sum_{m=1}^{2}\left(\mathbf{C}^{(m)} - \mathbf{B}^{(m)}\mathbf{F}^{(m)^{i\top}} - \mathbf{F}^{(m)^i}\mathbf{B}^{(m)\top} + \mathbf{F}^{(m)^i}\mathbf{A}^{(m)}\mathbf{F}^{(m)^{i\top}}\right) \tag{8}$$

which are essentially weighted average updates across the switching models.

Note that given knowledge about the structure in Eq 1, we may also further parameterize the state noise covariance matrix $\mathbf{Q}$ to be:

$$\mathbf{Q} = \begin{bmatrix} \sigma^2 & 0 \\ 0 & \sigma^2 \end{bmatrix} \quad \text{or} \quad \begin{bmatrix} \sigma_1^2 & 0 \\ 0 & \sigma_2^2 \end{bmatrix}$$

which can be readily accommodated by taking the trace or individual diagonal elements of the right hand sides of Eq 7 and Eq 8.

The state-transition matrix $\mathbf{F}^{(m)}$ needs to be handled differently since it has partial sharing of matrix elements. We can still derive the update equations by expanding out the elements of $\mathbf{A}^{(m)}$ and $\mathbf{B}^{(m)}$, resulting in row-wise update equations:

$$\begin{bmatrix} \mathbf{F}_{11}^{i} & \mathbf{F}_{12}^{(1)^i} \end{bmatrix} = \begin{bmatrix} \mathbf{B}_{11}^{(1)} + \mathbf{B}_{11}^{(2)} & \mathbf{B}_{12}^{(1)} \end{bmatrix} \left( \begin{bmatrix} \mathbf{A}_{11}^{(1)} + \mathbf{A}_{11}^{(2)} & \mathbf{A}_{12}^{(1)} \\ \mathbf{A}_{21}^{(1)} & \mathbf{A}_{22}^{(1)} \end{bmatrix} \right)^{-1} \tag{9}$$

$$\begin{bmatrix} \mathbf{F}_{21}^{i} & \mathbf{F}_{22}^{i} \end{bmatrix} = \begin{bmatrix} \mathbf{B}_{21}^{(1)} + \mathbf{B}_{21}^{(2)} & \mathbf{B}_{22}^{(1)} + \mathbf{B}_{22}^{(2)} \end{bmatrix} \left( \mathbf{A}^{(1)} + \mathbf{A}^{(2)} \right)^{-1} \tag{10}$$

which amount to partial sums of the hidden state moments across models. Then we have:

$$\mathbf{F}^{(1)^i} = \begin{bmatrix} \mathbf{F}_{11}^{i} & \mathbf{F}_{12}^{(1)} \\ \mathbf{F}_{21}^{i} & \mathbf{F}_{22}^{i} \end{bmatrix}, \qquad \mathbf{F}^{(2)^i} = \begin{bmatrix} \mathbf{F}_{11}^{i} & 0 \\ \mathbf{F}_{21}^{i} & \mathbf{F}_{22}^{i} \end{bmatrix}$$

In this simulation example, we fixed $\mathbf{G}$ to be the identity matrix. If needed, we can also update it:

$$\mathbf{G}^{i} = \left( \sum_{m=1}^{2} \sum_{t=1}^{T} h_t^{(m)} \mathbf{y}_t \mathbf{x}_{t|T}^{(m)\top} \right) \left( \sum_{m=1}^{2} \sum_{t=1}^{T} h_t^{(m)} \left( \mathbf{\Sigma}_{t|T}^{(m)} + \mathbf{x}_{t|T}^{(m)} \mathbf{x}_{t|T}^{(m)\top} \right) \right)^{-1} \tag{11}$$

Finally, the update equation for $\mathbf{R}$ is already provided in Materials and methods therefore omitted, and the update equations for the HMM process remain unchanged.

### Neural oscillations with nested structures

In the analyses of sleep oscillations, we assumed the following generative structure during learning:

$$\begin{aligned} \mathbf{x}_t^{(1)} &= \begin{bmatrix} a^{\delta}\mathcal{R}(\delta) & \mathbf{0} \\ \mathbf{0} & a^{\varsigma}\mathcal{R}(\varsigma) \end{bmatrix} \mathbf{x}_{t-1}^{(1)} + \mathbf{w}_t^{(1)}, \quad & \mathbf{w}_t^{(1)} &\sim \mathcal{N}\left( \mathbf{0}, \begin{bmatrix} (\sigma^2)^{\delta}\mathbf{I}_2 & 0 \\ 0 & (\sigma^2)^{\varsigma}\mathbf{I}_2 \end{bmatrix} \right) \\ \mathbf{x}_t^{(2)} &= a^{\delta}\mathcal{R}(\delta)\, \mathbf{x}_{t-1}^{(2)} + \mathbf{w}_t^{(2)}, & \mathbf{w}_t^{(2)} &\sim \mathcal{N}\left( \mathbf{0}, (\sigma^2)^{\delta}\mathbf{I}_2 \right) \end{aligned} \tag{12}$$

where $\mathbf{x}_t^{(1)} = \left[ x_{t,1}^{\delta(1)}, x_{t,2}^{\delta(1)}, x_{t,1}^{\varsigma(1)}, x_{t,2}^{\varsigma(1)} \right]^{\top}$ and $\mathbf{x}_t^{(2)} = \left[ x_{t,1}^{\delta(2)}, x_{t,2}^{\delta(2)} \right]^{\top}$;

$$\begin{aligned} y_t &= \mathbf{G}^{(s_t)} \mathbf{x}_t^{(s_t)} + v_t, \qquad v_t \sim \mathcal{N}(0, R) \\ \mathbf{G}^{(1)} &= \begin{bmatrix} 1 & 0 & 1 & 0 \end{bmatrix}, \quad \mathbf{G}^{(2)} = \begin{bmatrix} 1 & 0 \end{bmatrix} \end{aligned}$$

where $\mathbf{I}_2$ is an identity matrix, and for $f \in \{\delta, \varsigma\}$, $\mathcal{R}(f) = \begin{bmatrix} \cos\omega^f & -\sin\omega^f \\ \sin\omega^f & \cos\omega^f \end{bmatrix}$. The switching state $s_t$ follows an HMM process and takes on values in $\{1, 2\}$ across time points.

In this generative model, transient spindle activity is modelled by switching between two "neural" states where one has both slow oscillations ($\delta$) and spindles ($\varsigma$), while the other has only slow oscillations. By imposing the shared parameters $\left\{a^\delta, \mathcal{R}(\delta), (\sigma^2)^\delta\right\}$ between the slow oscillators in the two candidate models, this generative structure also captures the stationary dynamics of slow waves in the background regardless of the presence of spindles. To derive M-step update equations, we again write the expectation of the complete log-likelihood w.r.t. the approximate distribution $q$:

$$
\begin{aligned}
&\mathbb{E}\left[\log p\left(\left\{\mathbf{y}_t, \mathbf{x}_t^{(1)}, \mathbf{x}_t^{(2)}, s_t\right\} |\boldsymbol{\theta}\right)\right] \\
&= -\frac{T}{2}\log|2\pi\mathbf{R}| - \frac{1}{2}\sum_{m=1}^{2}\log\left|2\pi\mathbf{Q}_0^{(m)}\right| - \frac{T}{2}\sum_{m=1}^{2}\log\left|2\pi\mathbf{Q}^{(m)}\right| \\
&\quad - \frac{1}{2}\sum_{m=1}^{2}\sum_{t=1}^{T} h_t^{(m)}\operatorname{Tr}\left\{\mathbf{R}^{-1}\left(\left(\mathbf{y}_t - \mathbf{G}^{(m)}\mathbf{x}_{t|T}^{(m)}\right)\left(\mathbf{y}_t - \mathbf{G}^{(m)}\mathbf{x}_{t|T}^{(m)}\right)^\top + \mathbf{G}^{(m)}\boldsymbol{\Sigma}_{t|T}^{(m)}\mathbf{G}^{(m)\top}\right)\right\} \\
&\quad - \frac{1}{2}\sum_{m=1}^{2}\operatorname{Tr}\left\{\mathbf{Q}_0^{(m)-1}\left(\left(\mathbf{x}_{0|T}^{(m)} - \boldsymbol{\mu}\right)\left(\mathbf{x}_{0|T}^{(m)} - \boldsymbol{\mu}\right)^\top + \boldsymbol{\Sigma}_{0|T}^{(m)}\right)\right\} \\
&\quad - \frac{1}{2}\sum_{m=1}^{2}\operatorname{Tr}\left\{\mathbf{Q}^{(m)-1}\left(\mathbf{C}^{(m)} - \mathbf{B}^{(m)}\mathbf{F}^{(m)\top} - \mathbf{F}^{(m)}\mathbf{B}^{(m)\top} + \mathbf{F}^{(m)}\mathbf{A}^{(m)}\mathbf{F}^{(m)\top}\right)\right\} \\
&\quad + \sum_{m=1}^{2} p_{0|T}(m)\log\boldsymbol{\rho}_m + \sum_{t=1}^{T}\sum_{m,n=1}^{2} p_{t,t-1|T}(m,n)\log\boldsymbol{\phi}_{m,n}
\end{aligned}
\tag{13}
$$

Each oscillator is parameterized by $\boldsymbol{\theta}^f = \left\{a^f, \omega^f, (\sigma^2)^f, \boldsymbol{\mu}^f, (\sigma_0^2)^f\right\}$, $f \in \{\delta, \varsigma\}$. We can substitute them into Eq 13, which has a block-diagonal structure, and take individual partial derivatives to obtain the update equations. Detailed derivations of stationary oscillator update equations including Von Mises and inverse gamma priors are extensively described in [1], therefore omitted here.

At the $i$th EM iteration, the update equations for joint estimation of oscillator parameters corresponding to the slow oscillation component are listed here for reference:

$$
\boldsymbol{\mu}^{\delta^i} = \left[\boldsymbol{\mu}^i\right]_\delta, \qquad \text{where } \boldsymbol{\mu}^i = \frac{1}{2}\sum_{m=1}^{2}\mathbf{x}_{0|T}^{(m)} \tag{14}
$$

$$
(\sigma_0^2)^{\delta^i} = \frac{1}{4}\sum_{m=1}^{2}\operatorname{Tr}\left\{\left[\boldsymbol{\Sigma}_{0|T}^{(m)} + \mathbf{x}_{0|T}^{(m)}\mathbf{x}_{0|T}^{(m)\top} - \mathbf{x}_{0|T}^{(m)}\boldsymbol{\mu}^{i\top} - \boldsymbol{\mu}^i\mathbf{x}_{0|T}^{(m)\top} + \boldsymbol{\mu}^i\boldsymbol{\mu}^{i\top}\right]_\delta\right\} \tag{15}
$$

$$
\omega^{\delta^i} = \arctan\left(\frac{\operatorname{Rt}\left\{\mathbf{B}_\delta^{(1)} + \mathbf{B}_\delta^{(2)}\right\} + \kappa_{vm}^\delta\sin\left(\mu_{vm}^\delta\right)}{\operatorname{Tr}\left\{\mathbf{B}_\delta^{(1)} + \mathbf{B}_\delta^{(2)}\right\} + \kappa_{vm}^\delta\cos\left(\mu_{vm}^\delta\right)}\right) \tag{16}
$$

$$
a^{\delta^i} = \frac{\cos(\omega^{\delta^i})\operatorname{Tr}\left\{\mathbf{B}_\delta^{(1)} + \mathbf{B}_\delta^{(2)}\right\} + \sin(\omega^{\delta^i})\operatorname{Rt}\left\{\mathbf{B}_\delta^{(1)} + \mathbf{B}_\delta^{(2)}\right\}}{\operatorname{Tr}\left\{\mathbf{C}_\delta^{(1)} + \mathbf{C}_\delta^{(2)}\right\}} \tag{17}
$$

$$
(\sigma^2)^{\delta^i} = \frac{\beta_{ig}^\delta + \mathcal{S}/2}{a_{ig}^\delta + T/2 + 1} \tag{18}
$$

$$
\begin{aligned}
\text{where } \mathcal{S} = \frac{1}{4}\Big(&\operatorname{Tr}\left\{\mathbf{C}_\delta^{(1)} + \mathbf{C}_\delta^{(2)}\right\} \\
&- 2a^{\delta^i}\left(\cos(\omega^{\delta^i})\operatorname{Tr}\left\{\mathbf{B}_\delta^{(1)} + \mathbf{B}_\delta^{(2)}\right\} + \sin(\omega^{\delta^i})\operatorname{Rt}\left\{\mathbf{B}_\delta^{(1)} + \mathbf{B}_\delta^{(2)}\right\}\right) \\
&+ \left(a^{\delta^i}\right)^2\operatorname{Tr}\left\{\mathbf{A}_\delta^{(1)} + \mathbf{A}_\delta^{(2)}\right\}\Big)
\end{aligned}
$$

In the above set of equations, $[\mathbf{M}]_\delta$ (or simply $\mathbf{M}_\delta$) indicates the upper left block of a matrix $\mathbf{M}$. This block corresponds to the slow oscillation components in Eq 12. $\mathrm{Tr}\{\cdot\}$ is the matrix trace operator. Since all traces above are only taken on 2x2 matrices, then it is $\mathrm{Tr}\{\mathbf{M}\} = \mathbf{M}_{11} + \mathbf{M}_{22}$. $\mathrm{Rt}\{\cdot\}$ is a related operator on a 2x2 matrix that is $\mathrm{Rt}\{\mathbf{M}\} = \mathbf{M}_{21} - \mathbf{M}_{12}$. $\kappa^\delta_{vm}$ and $\mu^\delta_{vm}$ specify a Von Mises prior for the slow oscillation rotation frequency parameter $\omega^\delta$ as described in S2 Appendix. Likewise, $a^\delta_{ig}$ and $\beta^\delta_{ig}$ specify an inverse gamma prior for the state noise variance parameter $(\sigma^2)^\delta$.

We conclude by pointing out that joint estimation of parameters across nested models in this example again results in weighted averages and sums of hidden state estimates just like in Eq 6-10.

### Relations to alternative generative models

Eq 12 postulates two parallel Gaussian SSMs between which observations are obtained by selecting one of the two candidate models. Instead of this parallel model formulation, we can also consider a different generative model as follows:

$$\begin{bmatrix}\mathbf{x}^\delta_t\\ \mathbf{x}^\varsigma_t\end{bmatrix} = \begin{bmatrix}a^\delta\mathcal{R}(\delta) & \mathbf{0}\\ \mathbf{0} & a^\varsigma\mathcal{R}(\varsigma)\end{bmatrix}\begin{bmatrix}\mathbf{x}^\delta_{t-1}\\ \mathbf{x}^\varsigma_{t-1}\end{bmatrix} + \begin{bmatrix}\mathbf{w}^\delta_t\\ \mathbf{w}^\varsigma_t\end{bmatrix}, \quad \begin{bmatrix}\mathbf{w}^\delta_t\\ \mathbf{w}^\varsigma_t\end{bmatrix} \sim \mathcal{N}\left(\mathbf{0}, \begin{bmatrix}(\sigma^2)^\delta\mathbf{I}_2 & 0\\ 0 & (\sigma^2)^\varsigma\mathbf{I}_2\end{bmatrix}\right) \tag{19}$$

$$y_t = \mathbf{G}^{(s_t)}\begin{bmatrix}\mathbf{x}^\delta_t\\ \mathbf{x}^\varsigma_t\end{bmatrix} + v_t, \quad v_t \sim \mathcal{N}(0, R)$$
$$\mathbf{G}^{(1)} = \begin{bmatrix}1 & 0 & 1 & 0\end{bmatrix}, \quad \mathbf{G}^{(2)} = \begin{bmatrix}1 & 0 & 0 & 0\end{bmatrix}$$

As before, $\mathbf{I}_2$ is an identity matrix, and for $f \in \{\delta, \varsigma\}$:

$$\mathcal{R}(f) = \begin{bmatrix}\cos\omega^f & -\sin\omega^f\\ \sin\omega^f & \cos\omega^f\end{bmatrix},$$
$$\mathbf{x}^f_t = \begin{bmatrix}\mathbf{x}^f_{t,1}\\ \mathbf{x}^f_{t,2}\end{bmatrix}, \quad \mathbf{x}^f_{t-1} = \begin{bmatrix}\mathbf{x}^f_{t-1,1}\\ \mathbf{x}^f_{t-1,2}\end{bmatrix}, \quad \mathbf{w}^f_t = \begin{bmatrix}\mathbf{w}^f_{t,1}\\ \mathbf{w}^f_{t,2}\end{bmatrix}$$

This multi-modal formulation of slow oscillations and spindles assumes a single sequence of hidden states that evolve in the latent space. Transient spindle activity is then modeled after a switching observation matrix that either observes or ignores the spindle component. The generative process in Eq 19 ensures the smoothness of the slow oscillation component at the beginning and end of a spindle. In contrast, if we were to simulate sleep oscillations from Eq 12, there will be change-point discontinuities in the slow oscillation wave form around spindles.

Nevertheless, we emphasize that the assumed generative process in Eq 12 is a close approximation of Eq 19 for several reasons. First, Eq 12 imposes shared parameters for the slow oscillations across the two candidate models. Thus, the parameters encoding the dynamics of slow oscillations are the same across the parallel models, resulting in a single stationary model learned for the background slow waves. Second, hidden states are inferred from observed data during posterior inference. The parallel models in Eq 12 provide very similar smoothed estimates of slow oscillations after conditioning on the same observations. Therefore, the estimated slow oscillators in the two candidate models are not independent during switching inference. This is distinct from what would be two independent instances of the same model evolving separately when used to generate data, which results in potential discontinuities. Taken together, Eq 12 is effectively capturing the same time-varying dynamics as Eq 19 when employed for switching state-space modeling. But, Eq 12 can be solved directly using the variational learning framework presented in this paper with no modification of algorithmic steps (except joint estimation during M-steps, which have intuitive averaging forms as shown above). One can simply construct any numbers of parallel models capturing distinct neural dynamics of interest and use the exact same algorithm for switching inference. In contrast, generative structures like Eq 19 will require modified Kalman filtering and smoothing algorithms to cope with the changing observation matrices, as well as customized variational approximation of case-specific posterior distributions that are also intractable.

An implicit assumption made in both Eq 12 and Eq 19 is that there could be stationary spindle activity evolving in the hidden states, and temporal variations of spindles come from that the measured data do not observe them except transiently. To avoid this assumption, one could devise yet another generative process, which as we will show can again be closely approximated by Eq 12:

$$\begin{bmatrix}\mathbf{x}_t^{\delta}\\ \mathbf{x}_t^{\varsigma}\end{bmatrix} = \begin{bmatrix}a^{\delta}\mathcal{R}(\delta) & \mathbf{0}\\ \mathbf{0} & a^{\varsigma}\mathcal{R}(\varsigma)\end{bmatrix}\begin{bmatrix}\mathbf{x}_{t-1}^{\delta}\\ \mathbf{x}_{t-1}^{\varsigma}\end{bmatrix} + \begin{bmatrix}\mathbf{w}_t^{\delta}\\ \mathbf{w}_t^{\varsigma}\end{bmatrix}, \quad \begin{bmatrix}\mathbf{w}_t^{\delta}\\ \mathbf{w}_t^{\varsigma}\end{bmatrix} \sim \mathcal{N}\left(\mathbf{0}, \begin{bmatrix}(\sigma^2)^{\delta}\mathbf{I}_2 & 0\\ 0 & \mathbb{1}[s_t = 1](\sigma^2)^{\varsigma}\mathbf{I}_2\end{bmatrix}\right) \tag{20}$$

$$y_t = \begin{bmatrix}1 & 0 & 1 & 0\end{bmatrix}\begin{bmatrix}\mathbf{x}_t^{\delta}\\ \mathbf{x}_t^{\varsigma}\end{bmatrix} + v_t, \quad v_t \sim \mathcal{N}(0, R)$$

As before, $\mathbf{I}_2$ is an identity matrix, and for $f \in \{\delta, \varsigma\}$:

$$\mathcal{R}(f) = \begin{bmatrix}\cos\omega^f & -\sin\omega^f\\ \sin\omega^f & \cos\omega^f\end{bmatrix},$$

$$\mathbf{x}_t^f = \begin{bmatrix}\mathbf{x}_{t,1}^f\\ \mathbf{x}_{t,2}^f\end{bmatrix}, \quad \mathbf{x}_{t-1}^f = \begin{bmatrix}\mathbf{x}_{t-1,1}^f\\ \mathbf{x}_{t-1,2}^f\end{bmatrix}, \quad \mathbf{w}_t^f = \begin{bmatrix}\mathbf{w}_{t,1}^f\\ \mathbf{w}_{t,2}^f\end{bmatrix}$$

In this formulation, there is also a single multi-modal hidden state that evolves over time. But instead of a switching observation matrix, the state noise covariance for the spindle oscillator switches between $\mathbf{0}$ and $(\sigma^2)^{\varsigma}\mathbf{I}_2$. The key difference here is that the hidden spindle oscillator will have no activity when spindles are not present on the observed data, since the observation matrix is fixed and always emitting the spindle component. This might be more realistic when representing neural oscillations, as there is not an actual gating mechanism on measuring neural activity.

Our variational learning method on Eq 12 also recapitulates this behavior of Eq 20 during switching inference. During the fixed-point iterations in the E-step, Kalman smoothing (see Materials and methods and S2 Appendix) has modified observation noise that is inversely weighted by the model responsibility $h_t^{(m)}$. In other words, when the spindle component is unlikely to be present according to the switching state HMM, the smoothed estimates of the spindle component in the first model of Eq 12 are subject to large observation noise. This effectively skips the update equation during Kalman filtering and allows the spindle oscillator to quickly decay to zero during periods without spindles. In contrast, Eq 19 has a fixed observation noise and spindle state noise covariance. Therefore, if an inference algorithm is developed directly from Eq 19, it will take much longer for the spindle oscillator to decrease to zero, potentially not before the next spindle onsets.

Due to this feature of weighted Kalman smoothing, variational methods on Eq 12 allow parameter learning for the spindle oscillator while focusing on time points when spindles are statistically present. Direct inference on Eq 20 can also accomplish this, but it will create a singular state noise covariance matrix when spindles are not present, rendering standard Kalman filter, RTS smoother, and forward-backward algorithm inapplicable.

In summary, the parallel model formulation in Eq 12 gives a general solution to switching state-space modeling problems. It exhibits the same desired features in more carefully constructed multi-modal generative processes but conforms to efficient inference computations as described in this paper. We showed in this sleep oscillation example how Eq 12 can be viewed as close approximations to other generative structures in Eq 19 and Eq 20. Thus, these discussions highlight the effectiveness and generalizability of our variational learning method with joint estimation of parameters.
