## Supplementary material for "Switching state-space modeling of neural signal dynamics": S3 Appendix. Implementation details

### Forward-backward algorithm

To compute $h_t^{(m)}$ during the E-step and to obtain the posterior estimates of the discrete switching state, we need to follow a set of recursions known as the forward-backward algorithm [1] on the hidden Markov model (HMM) switching process. We utilize a normalized version of the algorithm to ensure numerical accuracy during time series analysis in practice.

An HMM is specified by a $M \times M$ state-transition matrix $\boldsymbol{\phi}$ and initial state probabilities $\boldsymbol{\rho}$. The algorithm begins with a forward pass:

$$\hat{\alpha}_t(m) = \frac{\exp\{g_t^{(m)}\} \sum_{s_{t-1}} \boldsymbol{\phi}_{m,s_{t-1}} \hat{\alpha}_{t-1}(s_{t-1})}{\sum_{s_t} \exp\{g_t^{(s_t)}\} \sum_{s_{t-1}} \boldsymbol{\phi}_{s_t,s_{t-1}} \hat{\alpha}_{t-1}(s_{t-1})} \tag{1}$$

with

$$\hat{\alpha}_0(m) = \boldsymbol{\rho}_m$$

Then the algorithm continues with a backward pass:

$$\hat{\beta}_t(m) = \frac{\sum_{s_{t+1}} \exp\{g_{t+1}^{(s_{t+1})}\} \boldsymbol{\phi}_{s_{t+1},m} \hat{\beta}_{t+1}(s_{t+1})}{\sum_{s_t} \sum_{s_{t+1}} \exp\{g_{t+1}^{(s_{t+1})}\} \boldsymbol{\phi}_{s_{t+1},s_t} \hat{\beta}_{t+1}(s_{t+1})} \tag{2}$$

with

$$\hat{\beta}_T(m) = 1, \; \forall\, m \in \{1, \cdots, M\}$$

Then the posterior smoothed estimates $h_t^{(m)} \triangleq \langle \mathbb{1}[s_t = m] \rangle_s$ can be computed as:

$$h_t^{(m)} = \frac{\hat{\alpha}_t(m) \hat{\beta}_t(m)}{\sum_{s_t} \hat{\alpha}_t(s_t) \hat{\beta}_t(s_t)} \tag{3}$$

To update the transition matrix $\boldsymbol{\phi}$ during the subsequent M-step, the posterior pairwise marginals are also needed. They can be easily computed as:

$$\langle \mathbb{1}[s_t = m, s_{t-1} = n] \rangle_s = \frac{\hat{\alpha}_{t-1}(n) \exp\{g_t^{(m)}\} \boldsymbol{\phi}_{m,n} \hat{\beta}_t(m)}{\sum_{s_t,s_{t-1}} \hat{\alpha}_{t-1}(s_{t-1}) \exp\{g_t^{(s_t)}\} \boldsymbol{\phi}_{s_t,s_{t-1}} \hat{\beta}_t(s_t)} \tag{4}$$

In S1 Appendix, we derived an expression for the negative variational free energy, which depends on calculating a normalizing constant from the HMM process using the forward pass variable of the forward-backward algorithm. As shown in [1], this normalization constant in the standard forward-backward algorithm can be calculated from the recursions without normalization:

$$\alpha_t(m) = \exp\{g_t^{(m)}\} \sum_{s_{t-1}} \boldsymbol{\phi}_{m,s_{t-1}} \alpha_{t-1}(s_{t-1})$$

$$\zeta = \sum_{m=1}^{M} \alpha_T(m) \tag{5}$$

In the normalized version outlined above, we forward pass the normalized forward variable $\hat{\alpha}_t(m)$ instead of $\alpha_t(m)$. Below, we derive a definition of $\zeta$ under normalization with proof by induction.

For ease of notation, we can denote the numerator in Eq 1 as $k_t(m)$:

$$k_t(m) \triangleq \exp\{g_t^{(m)}\} \sum_{s_{t-1}} \boldsymbol{\phi}_{m,s_{t-1}} \hat{\alpha}_{t-1}(s_{t-1})$$

therefore:

$$\hat{\alpha}_t(m) = \frac{k_t(m)}{\sum_{s_t} k_t(s_t)}$$

which together give the recursive relation:

$$k_t(m) = \exp\{g_t^{(m)}\} \sum_{s_{t-1}} \boldsymbol{\phi}_{m,s_{t-1}} \frac{k_{t-1}(s_{t-1})}{\sum_{s_{t-1}} k_{t-1}(s_{t-1})}$$

**Claim:** The normalization constant can be computed as:

$$\zeta = \prod_{t=1}^{T} \left( \sum_{m=1}^{M} k_t(m) \right) \tag{6}$$

**Proof:** Because the initial state probabilities $\boldsymbol{\rho}$ already sum to 1, the forward pass variable at $t = 0$ is the same regardless of normalization, i.e., $\hat{\alpha}_0(m) = \alpha_0(m)$. We then have at $t = 1$:

$$\begin{aligned} k_1(m) &= \exp\{g_1^{(m)}\} \sum_{s_0} \boldsymbol{\phi}_{m,s_0} \hat{\alpha}_0(s_0) \\ &= \exp\{g_1^{(m)}\} \sum_{s_0} \boldsymbol{\phi}_{m,s_0} \alpha_0(s_0) \\ &= \alpha_1(m) \end{aligned}$$

At the next time step $t = 2$ of the forward pass, we have:

$$\begin{aligned} k_2(m) &= \exp\{g_2^{(m)}\} \sum_{s_1} \boldsymbol{\phi}_{m,s_1} \frac{k_1(s_1)}{\sum_{s_1} k_1(s_1)} \\ &= \exp\{g_2^{(m)}\} \sum_{s_1} \boldsymbol{\phi}_{m,s_1} \alpha_1(s_1) \times \frac{1}{\sum_{s_1} k_1(s_1)} \\ &= \alpha_2(m) \times \frac{1}{\sum_{s_1} k_1(s_1)} \end{aligned}$$

By induction:

$$k_T(m) = \alpha_T(m) \prod_{t=1}^{T-1} \frac{1}{\sum_{s_t} k_t(s_t)} \tag{7}$$

Substituting Eq 7 into Eq 5 gives the expression in Eq 6 after simple rearrangement.

□

### de Jong smoothing

We utilize a set of recursions described in [2] to compute the posterior smoothed estimates of hidden states in Gaussian SSMs. This algorithm is analogous to Kalman filter followed by RTS smoother because it also has a forward and a backward pass. But it is more stable than the original Kalman smoothing by avoiding inversion of conditional state noise covariance, and it provides an efficient computation of interpolated densities that we use to initialize fixed-point iterations in the E-step.

For the $m^{\text{th}}$ Gaussian SSM, the algorithm begins with a forward pass:

$$\begin{aligned}
\mathbf{e}_t &= \mathbf{y}_t - \mathbf{G}\mathbf{x}_{t|t-1} \\
\mathbf{D}_t &= \mathbf{G}\boldsymbol{\Sigma}_{t|t-1}\mathbf{G}^\top + \mathbf{R}/h_t^{(m)} \\
\mathbf{K}_t &= \mathbf{F}\boldsymbol{\Sigma}_{t|t-1}\mathbf{G}^\top\mathbf{D}_t^{-1} \\
\mathbf{L}_t &= \mathbf{F} - \mathbf{K}_t\mathbf{G} \\
\mathbf{x}_{t+1|t} &= \mathbf{F}\mathbf{x}_{t|t-1} + \mathbf{K}_t\mathbf{e}_t \\
\boldsymbol{\Sigma}_{t+1|t} &= \mathbf{F}\boldsymbol{\Sigma}_{t|t-1}\mathbf{L}_t^\top + \mathbf{Q}
\end{aligned}$$

with the $^{(m)}$ superscripts dropped for all model parameters and with initial values

$$\begin{aligned}
\mathbf{x}_{1|0} &= \mathbf{F}\boldsymbol{\mu} \\
\boldsymbol{\Sigma}_{1|0} &= \mathbf{F}\mathbf{Q}_0\mathbf{F}^\top + \mathbf{Q}
\end{aligned}$$

Then a backward pass follows:

$$\begin{aligned}
\mathbf{r}_{t-1} &= \mathbf{G}^\top\mathbf{D}_t^{-1}\mathbf{e}_t + \mathbf{L}_t^\top\mathbf{r}_t \\
\mathbf{E}_{t-1} &= \mathbf{G}^\top\mathbf{D}_t^{-1}\mathbf{G} + \mathbf{L}_t^\top\mathbf{E}_t\mathbf{L}_t \\
\mathbf{x}_{t|T} &= \mathbf{x}_{t|t-1} + \boldsymbol{\Sigma}_{t|t-1}\mathbf{r}_{t-1} \\
\boldsymbol{\Sigma}_{t|T} &= \boldsymbol{\Sigma}_{t|t-1} - \boldsymbol{\Sigma}_{t|t-1}\mathbf{E}_{t-1}\boldsymbol{\Sigma}_{t|t-1} \\
\boldsymbol{\Sigma}_{t,t-1|T} &= \mathbf{L}_{t-1}\boldsymbol{\Sigma}_{t-1|t-2} - \boldsymbol{\Sigma}_{t|t-1}\mathbf{E}_{t-1}^\top\mathbf{L}_{t-1}\boldsymbol{\Sigma}_{t-1|t-2}
\end{aligned}$$

with the $^{(m)}$ superscripts also dropped for all smoothed estimates and with initial values

$$\mathbf{r}_T = \mathbf{0}, \;\; \mathbf{E}_T = \mathbf{0}$$

Our expression for the smoothed estimates of cross-covariance is novel and needs a simple proof.
**Proof:** Using [2, Theorem 1] and setting $s = t+1, m = t$:

$$\boldsymbol{\Sigma}_{t,s|T} = \boldsymbol{\Sigma}_{t,t+1|T} = \boldsymbol{\Sigma}_{t,t+1|t} - \boldsymbol{\Sigma}_{t|t-1}\mathbf{L}_t^\top\mathbf{E}_t\mathbf{L}_{t,t+1}\boldsymbol{\Sigma}_{t+1|t}$$

Then by [2, Lemma 1], we have:

$$\boldsymbol{\Sigma}_{t,t+1|t} = \boldsymbol{\Sigma}_{t|t-1}\mathbf{L}_t^\top, \;\; \mathbf{L}_{t,t+1} = \mathbf{I}$$

Therefore,

$$\boldsymbol{\Sigma}_{t,t+1|T} = \boldsymbol{\Sigma}_{t|t-1}\mathbf{L}_t^\top - \boldsymbol{\Sigma}_{t|t-1}\mathbf{L}_t^\top\mathbf{E}_t\boldsymbol{\Sigma}_{t+1|t}$$

By taking $t$ backward one time step and transposing the terms, We obtain the above expression.

□

During the backward recursions, most of the posterior estimates of hidden states can be computed exactly. However the intermediate variables $\mathbf{r}_t$ and $\mathbf{E}_t$ are not defined for $t = 0$. We can use the following approximations without significant impact to the variational learning algorithm:

$$\mathbf{x}_{0|T} = \mathbf{x}_{1|T}$$
$$\mathbf{\Sigma}_{1,0|T} = \mathbf{\Sigma}_{0|T} = \mathbf{\Sigma}_{1|T}$$

In S1 Appendix, we derived an expression for the negative variational free energy, which depends on calculating a normalization constant from the Gaussian SSMs using the marginal log-likelihood of observations. This can be computed in the innovations form using forward pass variables:

$$\log\ p(\{\mathbf{y}_t\}|) = -\frac{1}{2}\sum_{t=1}^{T}\log|2\pi\mathbf{D}_t| - \frac{1}{2}\sum_{t=1}^{T}\mathbf{e}_t^\top\mathbf{D}_t^{-1}\mathbf{e}_t$$

We use the term *interpolated density* to refer to the conditional density of $\mathbf{y}_t$ given all other observations, namely $\{\mathbf{y}_1, \cdots, \mathbf{y}_{t-1}, \mathbf{y}_{t+1}, \cdots, \mathbf{y}_T\}$. This density can be defined through the optimal linear projection of the random variable $\mathbf{y}_t$ onto the linear space spanned by the *leave-one-out* samples, $\{\mathbf{y}_1, \cdots, \mathbf{y}_{t-1}, \mathbf{y}_{t+1}, \cdots, \mathbf{y}_T\}$. Denoting this projection as $\mathbf{y}_{t|1,\cdots,t-1,t+1,\cdots,T}$, we consider the residual $\mathbf{y}_t - \mathbf{y}_{t|1,\cdots,t-1,t+1,\cdots,T}$ that follows a Gaussian distribution with [2, Theorem 4]:

$$\mathbf{y}_t - \mathbf{y}_{t|1,\cdots,t-1,t+1,\cdots T} = \mathbf{N}_t^{-1}\mathbf{n}_t$$
$$\mathbb{E}\left[\mathbf{y}_t - \mathbf{y}_{t|1,\cdots,t-1,t+1,\cdots T}\right] = \mathbf{0}$$
$$\mathbb{E}\left[\left(\mathbf{y}_t - \mathbf{y}_{t|1,\cdots,t-1,t+1,\cdots,T}\right)\left(\mathbf{y}_t - \mathbf{y}_{t|1,\cdots,t-1,t+1,\cdots,T}\right)^\top\right] = \mathbf{N}_t^{-1}$$

where $\mathbf{n}_t$ and $\mathbf{N}_t$ are defined in terms of the intermediate variables already computed during the forward and backward passes:

$$\mathbf{n}_t = \mathbf{D}_t^{-1}\mathbf{e}_t - \mathbf{K}_t^\top\mathbf{r}_t$$
$$\mathbf{N}_t = \mathbf{D}_t^{-1} + \mathbf{K}_t^\top\mathbf{E}_t\mathbf{K}_t.$$

Therefore the interpolated density can be expressed as:

$$p(\mathbf{y}_t|\mathbf{y}_{1,\cdots,t-1,t+1,\cdots,T}) = \frac{1}{\sqrt{|2\pi\mathbf{N}_t^{-1}|}}\exp\left\{-\frac{1}{2}\mathbf{n}_t^\top\mathbf{N}_t^{-1}\mathbf{n}_t\right\}.$$

### Iteration convergence

During the variational learning algorithm, there are two sets of iterations whose convergence need to be determined. First, there is the inner fixed-point iterations during each E-step. While an updated negative free energy can be computed after each round of updates of $h_t^{(m)}$ and $g_t^{(m)}$, it is often unnecessary and increases computational burden. Instead, we can check for convergence using the mean change of elements in the model evidence $g_t^{(m)}$. We stop the fixed-point iterations when this mean change of $g_t^{(m)}$ is below $10^{-6}$. Likewise, the outer EM iterations need to be checked for convergence. As analyzed in S1 Appendix, the negative free energy in this non-convex problem may decrease during consecutive EM iterations, therefore it is more reliable to check for empirical convergence using the mean change of elements in the converged model responsibility $h_t^{(m)}$ compared to the last E-step. We stop the EM iterations when this mean change of $h_t^{(m)}$ (converged in the E-step) is below $10^{-6}$.

### Model evidence in nested models

Candidate models in the oscillator simulation study have inherent nested structures, where the models with multiple oscillations encompass other models with subsets of those oscillations. Similarly in the spindle detection problem, transient spindles occur on top of an existing background activity of slow waves. This poses a new challenge for statistical inference on switching dynamics, which is fundamentally a problem of degrees of freedom: candidate models with more components have greater degrees of freedom to explain the temporal dynamics and will be favored in terms of log likelihood measures. It is important for switching inference methods to be able to handle nested models, because neural signals often involve overlaying transient activity on top of a stationary background, as exemplified by sleep spindles and slow oscillations in our example.

The current variational methods address this issue with the model responsibility $\mathbf{h}_t$ that weights the observed data during Kalman smoothing of the Gaussian SSMs (see the de Jong smoothing section above and Materials and methods). Once the fixed-point iterations have converged, the model evidence $g_t^{(m)}$ for each candidate model has been weighted by $h_t^{(m)}$. In other words, even though the model with more components among nested models will have "inflated" log likelihood values for all time points, its posterior model evidence will remain lower than the other candidate models on segments where the smoothed estimates from the HMM (i.e., $h_t^{(m)}$) suggest it to be less responsible for. It is therefore critical to initialize the fixed-point iterations during E-steps in an unbiased manner to account for this issue in nested models.

An adjustment is not needed for VI-A EM, as it already starts with equal model responsibilities $\mathbf{h}_t$ at a high temperature and anneals to stable values (albeit often inaccurate local maxima). One simple adjustment for VI-I EM is to mean-center the interpolated densities when used to initialize the model evidence $\mathbf{g}_t$ at every E-step. Using the spindle detection models as an example, Kalman smoothing of the two Gaussian SSMs produces the interpolated densities $p^{(1)}(y_t|y_{1,\cdots,t-1,t+1,\cdots,T})$ and $p^{(2)}(y_t|y_{1,\cdots,t-1,t+1,\cdots,T})$.

Then the variational model evidence $\mathbf{g}_t$ can be initialized as:

$$g_t^{(1)} = \log\left\{p^{(1)}(y_t|y_{1,\cdots,t-1,t+1,\cdots,T}) + \Delta^{(1)}\right\}$$
$$g_t^{(2)} = \log\left\{p^{(2)}(y_t|y_{1,\cdots,t-1,t+1,\cdots,T}) + \Delta^{(2)}\right\}$$

where

$$\Delta^{(1)} = \frac{\sum_{t=1}^{T} p^{(2)}(y_t|y_{1,\cdots,t-1,t+1,\cdots,T}) - \sum_{t=1}^{T} p^{(1)}(y_t|y_{1,\cdots,t-1,t+1,\cdots,T})}{2T}$$
$$\Delta^{(2)} = \frac{\sum_{t=1}^{T} p^{(1)}(y_t|y_{1,\cdots,t-1,t+1,\cdots,T}) - \sum_{t=1}^{T} p^{(2)}(y_t|y_{1,\cdots,t-1,t+1,\cdots,T})}{2T}$$

The above modification enables VI-I EM to remain sensitive to periods when the additional spindle component is absent. This technique extends to a set of $M$ candidate models in a straightforward manner by matching the ensemble averages of interpolated densities from different models.

### Priors for MAP estimates of sleep oscillations

Following [3], we added priors to the neural oscillator parameters to better constrain the EM learning of model parameters. We used a von Mises distribution, which is parameterized by a location measure $\mu_{vm}$ and a concentration measure $\kappa_{vm}$, on the rotation frequency parameter $\omega$:

$$p(\omega; \mu_{vm}, \kappa_{vm}) = \frac{1}{2\pi I_0(\kappa_{vm})} \exp\{\kappa_{vm} \cos(\omega - \mu_{vm})\}$$

where $I_0(\cdot)$ is the Bessel function of order 0 to make the density integrate to 1.

The location parameter $\mu_{vm}$ (in radian) was set to match 1 Hz and 13 Hz center frequencies, for slow oscillations and sleep spindles respectively, at the beginning of stationary Gaussian SSM learning. These modes updated to the learned oscillator frequencies before variational Bayesian learning. The concentration parameter $\kappa_{vm}$ was fixed to be high at 10,000.

We also added conjugate priors on the variance parameters $\left\{R, (\sigma^2)^\delta, (\sigma^2)^\varsigma\right\}$ using an inverse gamma distribution, which is parameterized by a shape parameter $\alpha_{ig}$ and a scale parameter $\beta_{ig}$:

$$p(\sigma^2; \alpha_{ig}, \beta_{ig}) = \frac{\beta_{ig}^{\alpha_{ig}}}{\Gamma(\alpha_{ig})}(1/\sigma^2)^{\alpha_{ig}+1} \exp\{-\beta_{ig}/\sigma^2\}$$

where $\Gamma(\cdot)$ is the gamma function. We fixed $\alpha_{ig}$ to be 10 and adjusted $\beta_{ig}$ so the modes of the inverse gamma prior were set to initial values at the onset of EM iterations. During the stationary Gaussian SSM learning, the modes for observation noise $R$ and state noise variances $\sigma^2$ for both slow oscillations and sleep spindles were set to be 1. For variational Bayesian learning, these modes were updated to the fitted parameter values after 50 EM iterations of the stationary Gaussian SSM.

The priors introduced here are not necessary for the variational learning algorithm to obtain reasonable spindle segmentation (results not shown). We added them to 1) show that MAP estimates can be easily incorporated in the current variational method, and to 2) help alleviate the non-convexity during learning to speed up convergence of the generalized EM iterations.
