## Supplementary material for "Switching state-space modeling of neural signal dynamics": S4 Appendix. Additional analyses

#### Segmentation with parameter learning

##### Negative free energy trajectories during EM iterations

Below we show two examples of negative free energy trajectory during generalized EM iterations from the simulation study of variational learning with two AR1 models. Variational learning with deterministic annealing (VI-A EM) and interpolated densities (VI-I EM) are compared, with each fixed to run 300 EM iterations regardless of convergence.

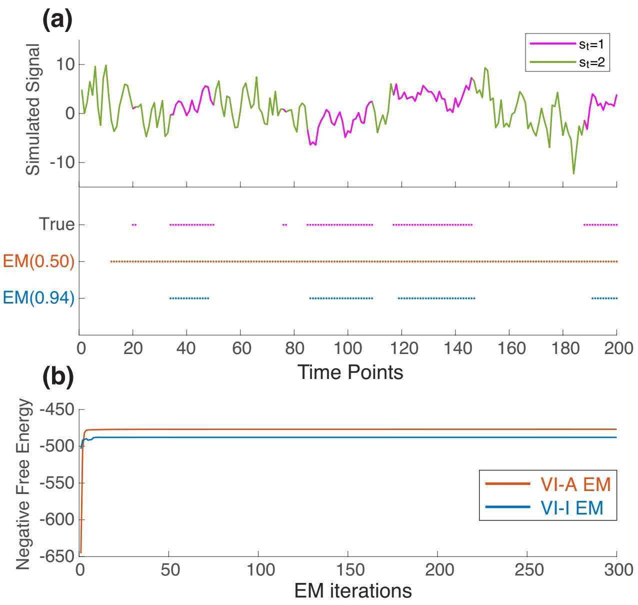


Figure S4-1: **Simulation results: negative free energy trajectories during EM iterations.** (a) A simulated sequence switching between two AR1 models with different dynamics. The top panel shows the time trace with two states marked in different colors. The bottom panel shows the segmentation results of variational EM learning on this example simulated data, where parameters are updated starting from randomly initialized values. Time points estimated to be in the first candidate model ($s_t = 1$) are marked in colored dots for both variational learning methods, with accuracy displayed in parentheses. (b) Negative free energy trajectories for the two variational learning methods. The plotted negative free energy values were computed after each E-step reached convergence. VI-A EM = variational EM learning with deterministic annealing (orange color); VI-I EM = variational EM learning with interpolated densities (blue color).

The first example in Fig S4-1 shows a case where the VI-A EM resulted in a trivial segmentation, labelling all time points as from the second candidate model with larger state noise variance.

Inspecting the negative free energy trajectories shows that absolute values of negative free energy alone do not provide a sufficient performance comparison between learning algorithms. Despite having a higher bound on the marginal log-likelihood of observations, VI-A EM had much worse segmentation accuracy than VI-I EM. This is because VI-A EM got trapped in a local maximum and produced degenerate parameters and $q$ estimates that ignored dynamic switching.

The second example in Fig S4-2a shows a different case where the VI-A EM method produced more reasonable segmentation results. In this case, the negative variational free energy, which gives a lower bound on the marginal log-likelihood of observations, is higher for VI-I EM than for VI-A EM. This is in agreement with the better segmentation accuracy obtained by the VI-I EM method.

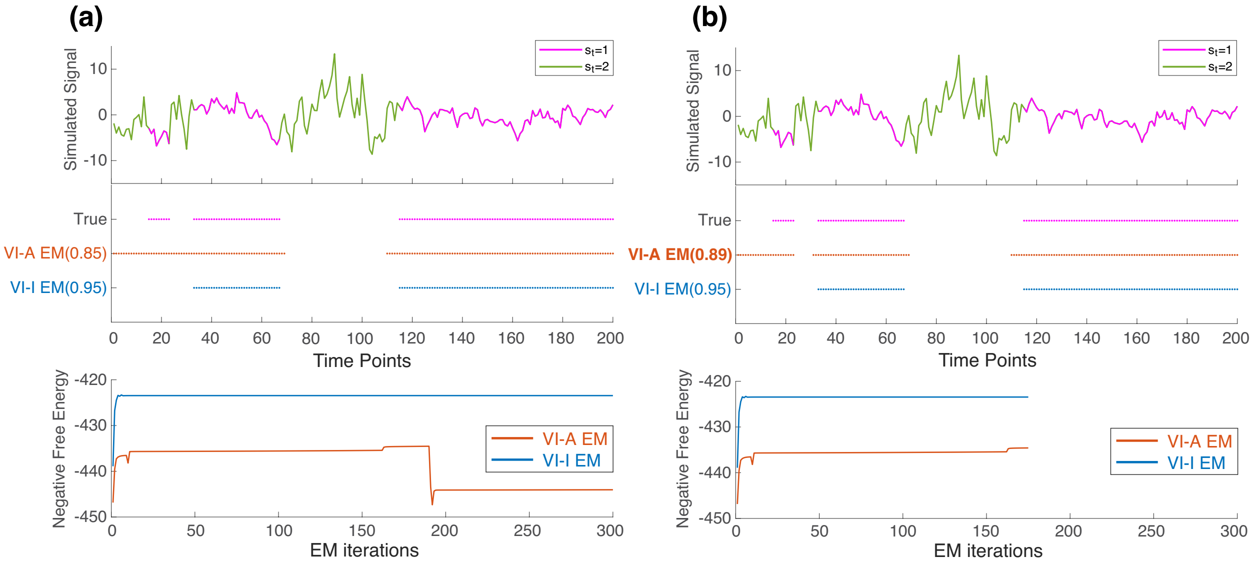


Figure S4-2: **Simulation results: negative free energy trajectories during EM iterations (Continued).** (a) Another simulated sequence switching between two AR1 models with different dynamics. The top panel shows the time trace with the two states, and the middle panel shows the segmentation results of variational EM learning. Time points estimated to be in the first candidate model ($s_t = 1$) are marked in colored dots, with accuracy displayed in parentheses. The bottom panel displays the negative free energy values, computed after each E-step convergence. (b) Variational learning on the same sequence with the EM algorithms stopped after 175 iterations. The segmentation was better for VI-A EM; correspondingly, the negative free energy trajectory ended at a higher value as shown in the bottom panel. In contrast, the segmentation accuracy and negative free energy values have already stabilized for VI-I EM and do not differ from those obtained after 300 iterations. VI-A EM = variational EM learning with deterministic annealing (orange color); VI-I EM = variational EM learning with interpolated densities (blue color).

Both examples also illustrate that the negative free energy trajectories could occasionally decrease during the EM iterations. This appears to contradict the theoretical motivation of generalized EM algorithm that should result in non-decreasing trends of negative free energy optimization. This discrepancy can be reconciled by noting that in this non-convex problem, the fixed-point iterations during the E-step only reach local maxima rather than the global maximum. In addition, both deterministic annealing and interpolated densities alter the starting points of the E-step after parameter updates from the last M-step. Therefore, it is possible that a shallower local negative free energy maximum, which has higher KL divergence from the true posterior, is reached during the next fixed point iteration. When this occurs, the empirically computed negative free energy could suddenly decrease, as seen for VI-A EM towards the end of iterations in Fig S4-2a. A truncated EM after 175 iterations as shown in Fig S4-2b demonstrates that the large decrease in negative free energy beyond 200 EM iterations indeed corresponded to a reduced segmentation accuracy than what was already achieved early on.

These two examples make it clear that the advantage of using interpolated densities lies in E-step initialization that is closer to a reasonable switching segmentation optimum. Interpolated densities do so by statistically comparing the candidate models at every time point. Such informed initialization technique to some extent avoids close-by local maxima that could sway the learning path when an approach like deterministic annealing is taken. With a slightly shallower local maximum at the end of the fixed-point iterations, VI-A EM often suffers consecutive decreases in the negative free energy, which is less frequent for VI-I EM. This behavior is reflected by the small dips in the negative free energy trajectories during early iterations shown in Fig S4-1 and Fig S4-2.

#### Relative sensitivity of parameter updates

In this simulation study, the observation noise variance $R$ did not get updated substantially (i.e., remained near the initial values) throughout EM iterations in both VI-A EM and VI-I EM. This resulted in a converged distribution similar to the uniform distribution used for initializing $R$, unlike the transition matrix $F$ or state noise variance $Q$ that showed distributional modes at the true values for VI-I EM. This relative insensitivity of $R$ during EM iterations stems from a much smaller magnitude compared to $Q$. Thus, the negative variational free energy profile against $R$ is flatter (locally) than against $Q$. To demonstrate this, we visualized negative variational free energy profile against $R$ and $Q$ with other parameters fixed at true values, respectively in Fig S4-3.

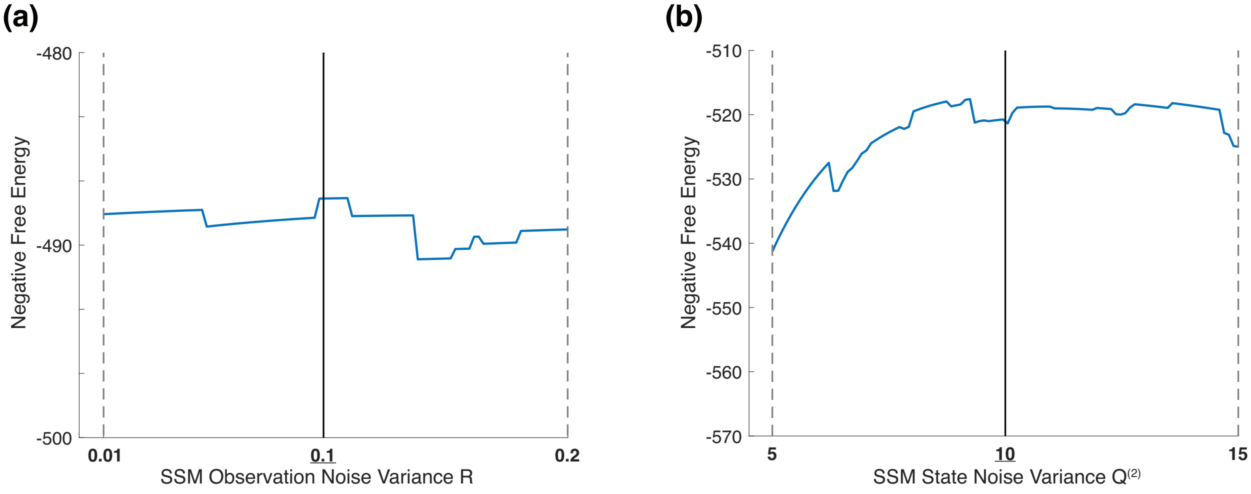


Figure S4-3: **Negative variational free energy as a function of $R$ and $Q$ for variational inference with interpolated densities.** (a) A plot of negative free energy as the value of $R$ varies across the initial distribution range, which shows a flattened curve around the true value at 0.1. (b) A plot of negative free energy as the value of $Q^{(2)}$ varies across the initial distribution range, which peaks near the true value at 10 with curvature around it, displaying increased sensitivity compared to varying the value of $R$.

#### Initialize parameters with less informative distributions

Although the initial parameters used in this simulation study were randomly drawn from uniform distributions, these distributions were centered around true parameters and potentially provide information that could facilitate switching segmentation. In many neuroscience applications, model parameters can indeed be reasonably initialized such as in the real-world spindle detection example where stationary models can be used to estimate parameters of Gaussian SSMs in a data-driven way. Nevertheless, here we repeat our simulation study with univariate AR1 models after initializing the model parameters using less informative distributions. We analyzed the segmentation accuracy and parameter recovery performance as in the main study. Specifically, we initialized all algorithms with random parameters drawn from uniform distributions for the $l^{\text{th}}$ sequence as following:

$$\begin{aligned}
F^{(1)} &= 0.90, & F_l^{(1)} &\sim \mathcal{U}_{[0.6,1.0]} \\
F^{(2)} &= 0.70, & F_l^{(2)} &\sim \mathcal{U}_{[0.6,1.0]} \\
Q^{(1)} &= 2, & Q_l^{(1)} &\sim \mathcal{U}_{[1,15]} \\
Q^{(2)} &= 10, & Q_l^{(2)} &\sim \mathcal{U}_{[1,15]} \\
R &= 0.1, & R_l &\sim \mathcal{U}_{[0.01,0.2]} \\
\phi_{11} = \phi_{22} &= 0.95, & \phi_{11,l} = \phi_{22,l} &\sim \mathcal{U}_{[0.9,0.99]}
\end{aligned}$$

Since the same distributions with wider supports are used to sample transition matrix $F$ and state noise variance $Q$ values for the two switching models, there would be an ordering ambiguity in labelling the learned models and switching states. To resolve this permutation issue, we post-hoc labelled the model with smaller estimated $Q$ as the first model, and the other as the second model, since the biggest difference between the two models is the magnitude of $Q$.

Results of this analysis replicate the patterns obtained when using informative distributions: despite overall reduced segmentation accuracy, VI-I EM outperformed the other methods (Fig S4-4a). Consistent with the wider support of initial value distributions, the converged parameter estimates in Fig S4-4b have wider spread. But VI-I EM again achieved better parameter recovery than VI-A EM by skewing more towards the true values. Notably, the segmentation accuracy of VI-I EM increased linearly against a geometric increase (by a factor of 2) in the data length, while the accuracy of other methods stayed constant and lower in comparison (Fig S4-4c). Lastly, parameter updates took more EM iterations to converge and on average incurred larger estimation errors compared to when informative distributions were used (Fig S4-4d). Nevertheless, VI-I EM showed robust convergence of parameter estimates towards the true values, which improved further with a longer data length.

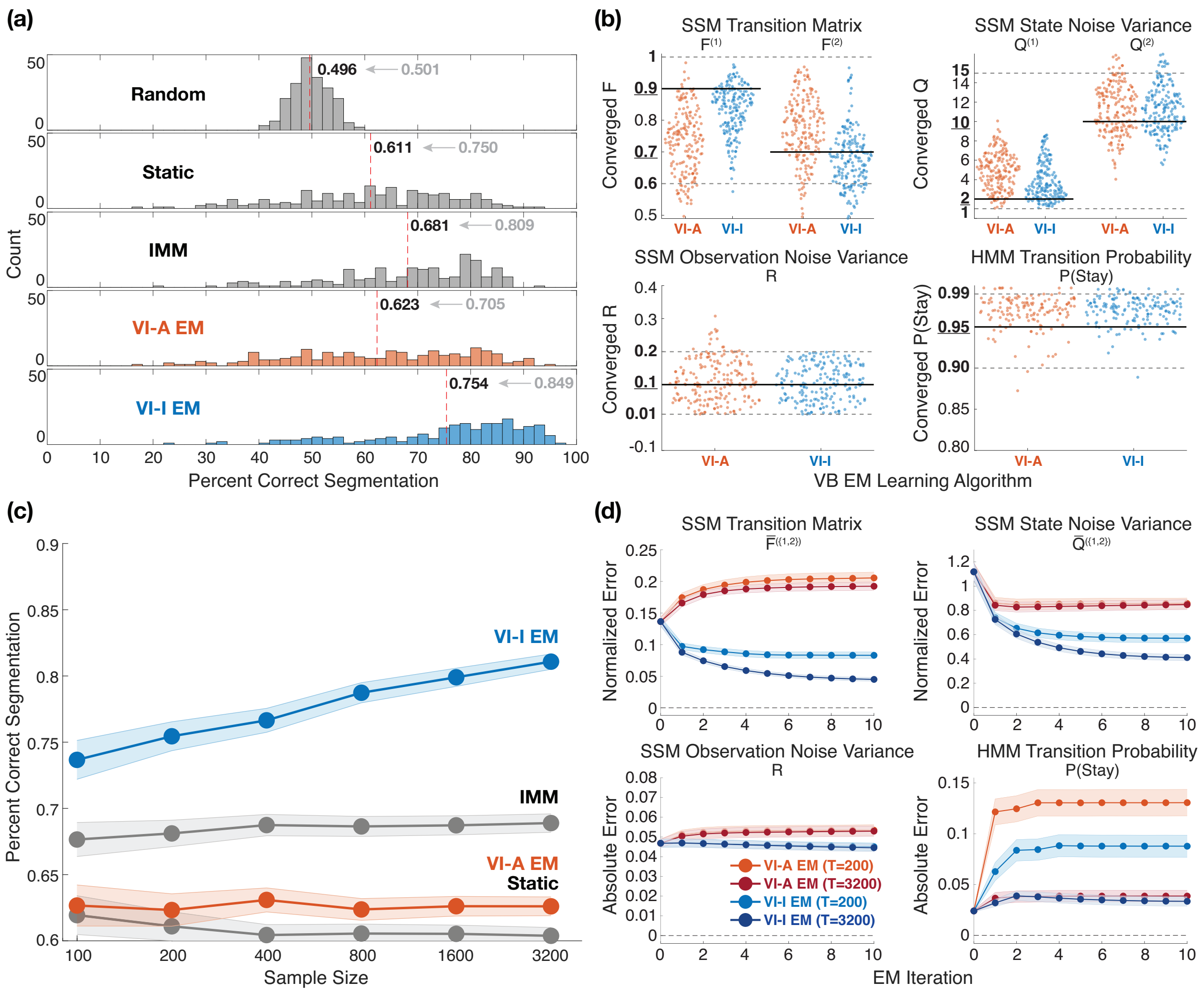


Figure S4-4: **Simulation results: segmentation performance for less informative initial value distributions.** (a) Histograms of segmentation accuracy with parameter learning across 200 repetitions of sequences of 200 time points. The mean segmentation accuracy for each method is displayed and marked by the dashed red line. The mean accuracy previously obtained using informative distributions is shown in gray. Random = random segmentation with a Bernoulli process; Static = static switching method; IMM = interacting multiple models method; VI-A EM = variational EM learning with deterministic annealing (orange color); VI-I EM = variational EM learning with interpolated densities (blue color). (b) Swarm plots showing the distributions of model parameters learned by the variational learning algorithms for sequences of 200 time points. Uniform distributions used to sample initial parameters are marked in bold fonts on the y-axes, as well as using solid black lines (true values) and dotted gray lines (upper and lower bounds of the ranges). (c) Changes of mean segmentation accuracy over sequences of varying data lengths. Shaded bounds denote the standard error of the mean around the average accuracy values. (d) Mean parameter estimation errors from true values across 10 EM iterations for two different data lengths. For transition matrix $F$ and state noise variance $Q$, normalized error is defined as abs(estimated - true)/true, and averaged across the two switching models. Absolute error is defined as abs(estimated - true). Shaded bounds denote the standard error of the mean around the average error values.

### Extensions of model structure and parameter estimation

#### Distributions of estimated model parameters

Here we show the distribution plots of estimated model parameter values in the bivariate AR1 simulations. As can be observed on Fig S4-5, VI-I EM and VI-A EM achieved comparable parameter recovery. Distributional modes are centered at the true values for the transition matrix $\mathbf{F}$ and state noise variances $\mathbf{Q}$. The same insensitivity of the observation noise variance $\mathbf{R}$ to EM updates as in the previous simulation study was encountered here, likely due to the small magnitude of $\mathbf{R}$ compared to that of $\mathbf{Q}$. Similarly, the bias in estimating the probability of staying in the same model for the HMM switching process is present due to the short data length of 200 time points.

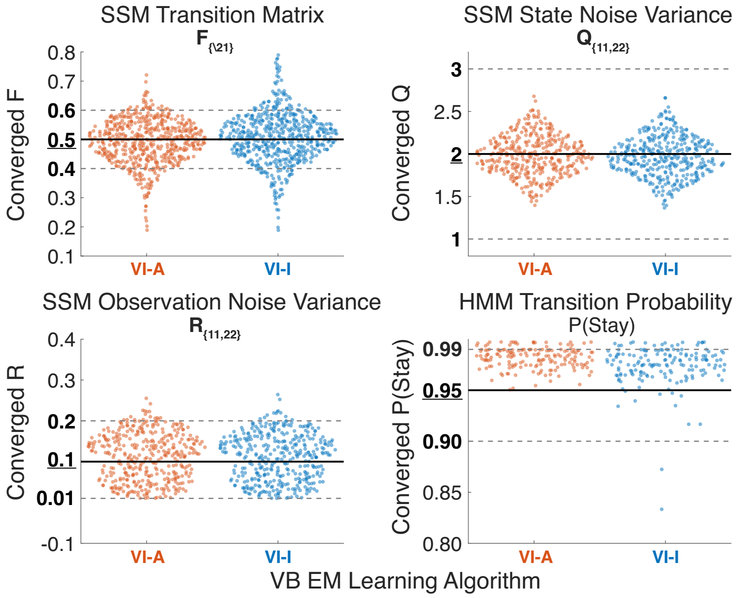


Figure S4-5: **Distributions of learned model parameters in the bivariate AR1 simulations.** Swarm plots showing the distributions of model parameters learned by the variational learning algorithms. Uniform distributions used to sample initial parameters are marked in bold fonts on the y-axes, as well as using solid black lines (true values) and dotted gray lines (upper and lower bounds of the ranges). Since the matrix elements of $\mathbf{F}$, $\mathbf{Q}$, and $\mathbf{R}$ have the same initial value distributions and true values and are shared across the two switching states, estimated entries of each parameter matrix are plotted on the same swarm plot, resulting in 600 points for $\mathbf{F}_{\{\backslash 21\}}$ and 400 points for $\mathbf{Q}$ and $\mathbf{R}$ from 200 repetitions. VI-A EM = variational EM learning with deterministic annealing (orange color); VI-I EM = variational EM learning with interpolated densities (blue color).

#### Comparisons to variational algorithms with an accurate generative model

As noted in the main Results section, our variational learning algorithm assumes a different generative model from the actual generative process underlying the bivariate AR1 simulations.

The true generative model is as following:

$$\mathbf{x}_t = \begin{bmatrix} 0.5 & \mathbf{F}_{12}^{(s_t)} \\ 0 & 0.5 \end{bmatrix} \mathbf{x}_{t-1} + \mathbf{w}_t, \qquad \mathbf{w}_t \sim \mathcal{N}\left(\mathbf{0}, \begin{bmatrix} 2 & 0 \\ 0 & 2 \end{bmatrix}\right)$$

$$\mathbf{y}_t = \begin{bmatrix} 1 & 0 \\ 0 & 1 \end{bmatrix} \mathbf{x}_t + \mathbf{v}_t, \qquad \mathbf{v}_t \sim \mathcal{N}\left(\mathbf{0}, \begin{bmatrix} 0.1 & 0 \\ 0 & 0.1 \end{bmatrix}\right)$$

where $\mathbf{F}_{12}^{(1)} = 0.5$ and $\mathbf{F}_{12}^{(2)} = 0$. This is a process with a single bivariate hidden state and a switching model parameter. Such generative model structure has been directly considered in some recent variational inference and learning algorithms for switching state-space models [1, 2, 3]. To better benchmark the performance of segmentation accuracy of VI-I EM, we applied three of such algorithms available at https://github.com/lindermanlab/ssm, which assume the accurate generative model structure, on the same bivariate AR1 simulations in this study:

1. Black-box stochastic variational inference with mean-field approximation
2. Black-box stochastic variational inference with tridiagonal Hessian
3. Laplace-EM with structured posterior as the approximate distribution $q$ in VI-I EM

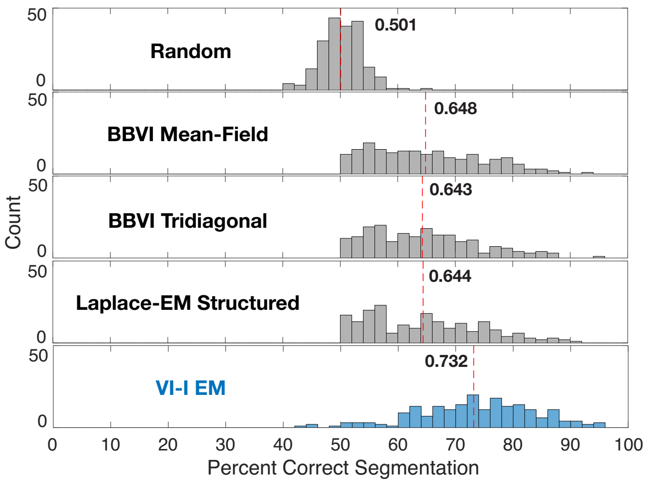


Figure S4-6: **Simulation results: segmentation performance using different variational algorithms in the bivariate AR1 simulations.** Histograms of segmentation accuracy across 200 repetitions. The mean segmentation accuracy for each method is displayed and marked by the dashed red line. BBVI = black-box variational inference; VI-I EM = variational EM learning with interpolated densities (blue color).

Fig S4-6 shows that VI-I EM is able to achieve superior segmentation accuracy compared to algorithms that assume the correct generative model structure. However, it should be noted that the three variational algorithms tested here used AR-HMM to initialize the model parameters from scratch, which performed better than specifying the model parameter at the true values (results not shown). In comparison, VI-I EM relied on initial model parameters that were informative of the true values, but had the advantage of utilizing a more specified model structure. In addition, the use of exact message passing and parameter updates along with interpolated densities in VI-I EM likely better exploits prior knowledge of the switching process. In essence, VI-I EM requires more informative candidate model construction compared to these other variational algorithms with more flexible approximations, resulting in better segmentation performance.

### Real-world application: spindle detection

#### Soft and hard segmentation with VI-I EM

As noted in the Materials and methods section, the estimated posterior model probabilities of VI-A and VI-I are polarized to take values close to either 1 or 0 with very sharp transitions. However in some applications, we may prefer a softer segmentation with gradual transitions to reflect the uncertainty around the transition periods in between two switching states. In other cases, discrete labels on the posterior switching states may be desired. To address these demands, we propose the following techniques to achieve soft and hard segmentation respectively for our variational switching inference method:

1. **Soft segmentation:** Once VI-I EM learning has converged, updated model parameters are available for all candidate models and the HMM process. Using the learned parameters, we first compute the interpolated densities at each time point for each candidate model and run the forward-backward algorithm on the HMM process using the interpolated densities as the observation probability. Then an approximate soft segmentation can be achieved using the smoothed probabilities of the discrete switching states in HMM taking each possible value.

2. **Hard segmentation:** This could be obtained by thresholding the polarized posterior model probabilities in VI-I but may not achieve the most likely sequence for the entire time series. Instead we again compute the interpolated densities at each time point for each candidate model and then run the Viterbi algorithm on the HMM process with the interpolated densities as the observation probability to identify the most likely sequence of the discrete states.

Since interpolated densities are integral to both approximate segmentation techniques, normalization for model complexity is necessary for nested structures. We applied the soft and hard segmentation to the short EEG segment analyzed for spindle detection. Fig S4-7 shows that both soft and hard segmentation agree well with the VI-I EM posterior model probabilities. The soft segmentation has the desired gradual transitions around sleep spindles and crosses 0.5 around the margins identified by VI-I EM. The hard segmentation is almost identical to the VI-I EM posterior model probabilities with slight differences on the boundaries of spindles, overall resulting in spindles of slightly shorter duration than those segmented by VI-I EM.

The soft and hard segmentation methods here can be viewed as generalizations of the static method in [4]. The static method uses the predicted density $p(y_t|x^{(m)}_{t|t-1})$ as a surrogate for the predicted observation probability $p(y_t|s_t = m, y_1, \cdots, y_{t-1})$ in an HMM. This is an approximation because the actual density $p(y_t|s_t = m, y_1, \cdots, y_{t-1})$ requires marginalizing out the entire history for $\{s_{t-1}, \cdots, s_1\}$ when computing $p(y_t|x^{(m)}_{t|t-1})$. In the same vein of approximation and Gaussian merging, we are approximating the smoothed density $p(y_t|s_t = m, y_1, \cdots, y_{t-1}, y_{t+1}, \cdots, y_T)$ with the interpolated density from each candidate model and running the forward-backward algorithm to obtain smoothed estimates of the switching state. Due to the approximations involved, it is difficult to quantify the validity of the soft and hard segmentation methods other than checking the segmentation accuracy. However, in the case of the soft segmentation, the segmentation itself may not be of interest to researchers since the proposed VI-I EM can achieve that already. Instead, the usefulness of the soft segmentation might have to be assessed empirically within a specific application. The soft segmentation method emphasizes the transition periods between states rather than multiple segments of a time series with distinct dynamics. Thus, its usage is more relevant in applications where key hypotheses are focused on the transition periods between segments.

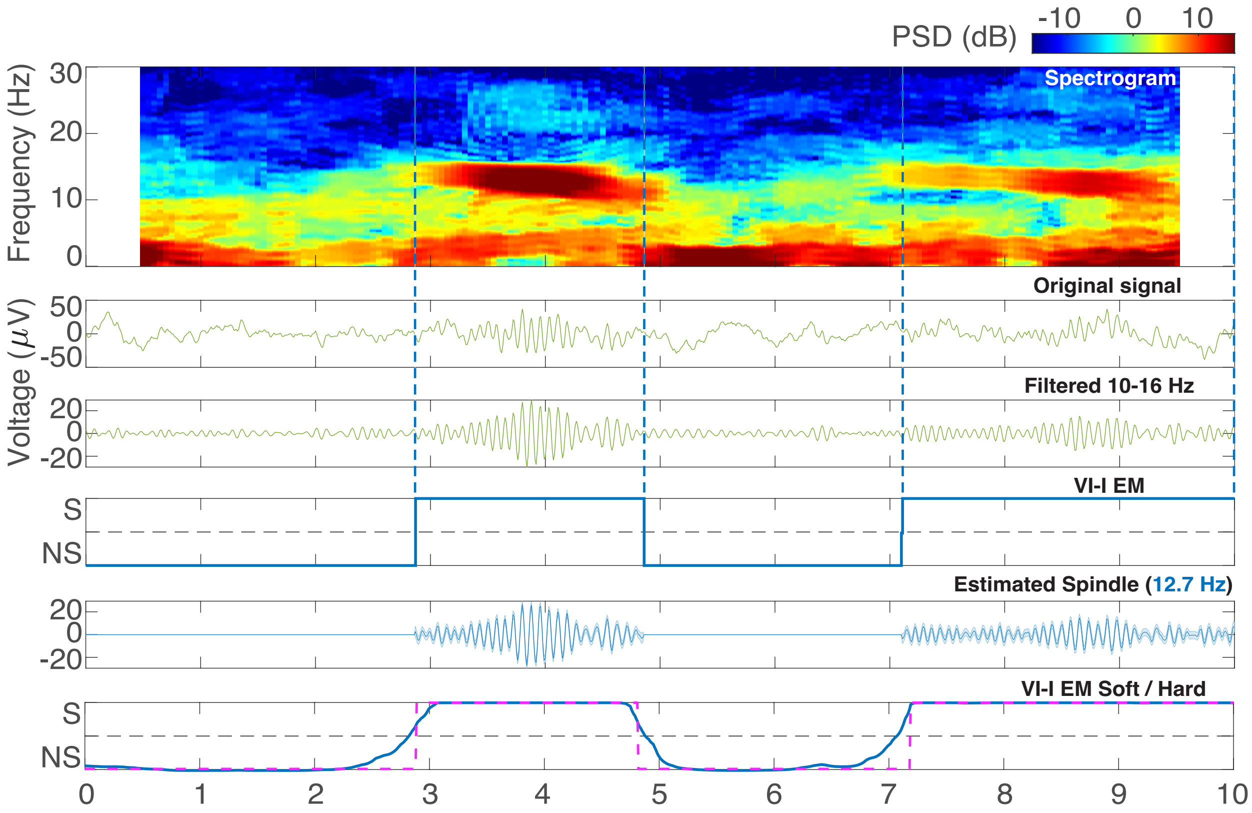


Figure S4-7: **Spindle detection with soft and hard segmentation.** In the top three panels, the spindle activity is visualized using a spectrogram, the original time trace, and after being bandpass filtered within 10 Hz to 16 Hz. The fourth panel shows the posterior model probabilities of $s_t = 1$ estimated by variational EM learning with interpolated densities (VI-I EM, blue color). The margins of spindle events identified by VI-I EM are also marked with vertical dashed lines. The fifth panel displays the estimated real part of spindle waveform with 95% confidence intervals according to posterior covariances. The learned spindle center frequency is displayed in blue in parentheses. The last panel shows the posterior switching state estimates under soft and hard segmentation. Soft estimates are smoothed estimates on the HMM process (blue solid line), while hard estimates are the most likely sequence from the Viterbi algorithm on the HMM process (magenta dashed line).

#### Real-time detection of sleep spindles

With the switching state-space modeling framework proved to be useful for detecting spindles, a related question is whether this model can be used to detect spindles in real-time as required in certain applications such as optogenetic modulation in animal studies or acoustic stimulation in human sleep studies. Given the generative model, the inference algorithm can be adapted to label new observations as they arrive as presence or absence of spindles in real time. Specifically, we will require the filtered estimate of the posterior model probabilities, which can be easily computed by performing variational inference (i.e., without M-step) on a new chunk of data with a given set of learned model parameters.

We applied this method on one of the 30 s NREM-2 sleep EEG recordings analyzed for spindle detection. We first ran the VI-I EM to convergence to obtain the model parameters, then we performed variational inference on a moving window of 10 s and recorded the filtered estimate of the model probability at the last sample of each window. Fig S4-8 shows the result of such real-time detection of sleep spindles in comparison to the usual smoothed posterior estimates by VI-I EM. Many false positives are present with very short duration, likely a result of window edge effects of variational inference on isolated abrupt changes in the time trace. Nevertheless, the method is able to flag a spindle consistently with about 100 ms delay.

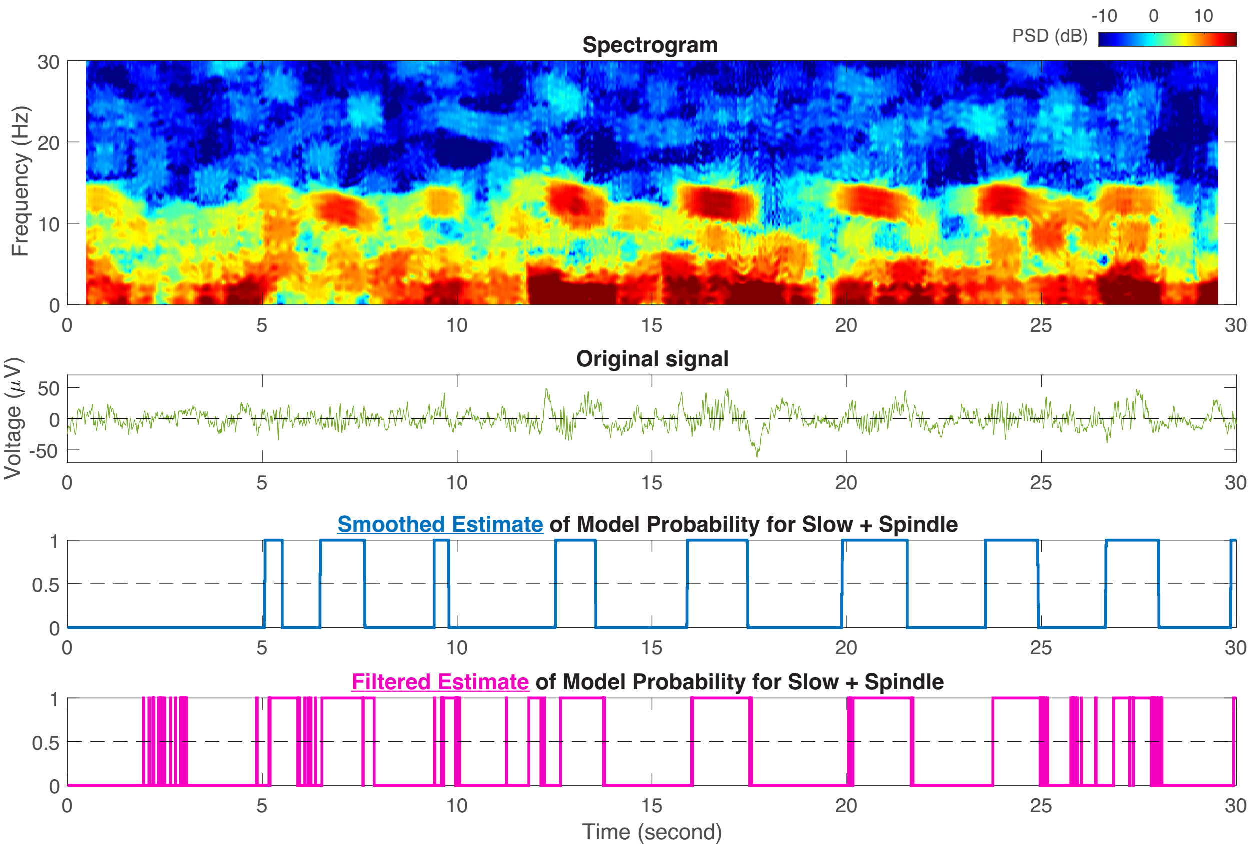


Figure S4-8: **Real-time detection of sleep spindles with variational inference.** The spindle activity is visualized using a spectrogram and the original time trace in the top two panels. The last two panels show smoothed (blue) and filtered (magenta) estimates of the posterior model probabilities of $s_t = 1$ estimated by variational inference with interpolated densities.
